# Infection-induced miR-126 suppresses *tsc1*- and *cxcl12a*-dependent permissive macrophages during mycobacterial infection

**DOI:** 10.1101/2021.10.07.463594

**Authors:** Kathryn Wright, Kumudika de Silva, Karren M. Plain, Auriol C. Purdie, Warwick J. Britton, Stefan H. Oehlers

**Affiliations:** Tuberculosis Research Program at the Centenary Institute, The University of Sydney, Camperdown NSW Australia; The University of Sydney, Faculty of Science, Sydney School of Veterinary Science, Sydney NSW, Australia; Department of Clinical Immunology, Royal Prince Alfred Hospital, Camperdown, NSW, Australia; The University of Sydney, Faculty of Medicine and Health & Sydney Institute for Infectious Diseases, Camperdown NSW Australia

**Author notes:** Corresponding author: Stefan Oehlers, Postal address: Centenary Institute, Locked Bag 6, Newtown, NSW 2042, Australia.

## Abstract

Regulation of host microRNA (miRNA) expression is a contested node that controls the host immune response to mycobacterial infection. The host must overcome concerted subversive efforts of pathogenic mycobacteria to launch and maintain a protective immune response. Here we examine the role of miR-126 in the zebrafish model of *Mycobacterium marinum* infection and identify a protective role for this infection-induced miRNA through multiple effector pathways. Specifically, we analyse the impact of the miR-126 knockdown-induced *tsc1a* and *cxcl12a/ccl2/ccr2* signalling axes during early host-*M*. *marinum* interactions. We find a strong detrimental effect of *tsc1a* upregulation that renders zebrafish embryos susceptible to higher bacterial burden and increased cell death despite dramatically higher recruitment of macrophages to the site of infection. We demonstrate that infection-induced miR-126 suppresses *tsc1* and *cxcl12a* expression thus improving macrophage function early in infection, partially through activation of mTOR signalling and strongly through preventing the recruitment of Ccr2+ permissive macrophages, resulting in the recruitment of protective *tnfa*-expressing macrophages. Together our results demonstrate an important role for infection-induced miR-126 in shaping an effective immune response to *M*. *marinum* infection in zebrafish embryos.

## Introduction

Manipulation and subversion of host signalling pathways is a hallmark of infection by pathogenic mycobacteria, such as the causative agents of tuberculosis and leprosy ^1,2^. Infection with pathogenic mycobacteria results in dysregulated immune responses and disruption of key signalling cascades, ultimately supporting the survival and persistence of bacteria ^3^. A crucial element of mycobacterial persistence is the interaction between bacteria and innate immune cells, primarily macrophages. Subversion of this normally protective interaction between macrophage and bacterium leads to the formation of granulomas which aid bacterial survival. Pleiotropic modulation of host gene expression and signalling cascades by microRNA (miRNA) potentially act as master regulators of the immune cell response to mycobacteria and may shape the outcome of infection.

miRNA are short single-stranded non-coding RNA molecules that post-transcriptionally regulate gene expression. Through a process known as gene-silencing, miRNA bind to the untranslated regions (UTRs) of target mRNA and reduce their stability. Depending on the degree of complementarity in base pairing, miRNA may either degrade the bound mRNA, or transiently suppress translation, reducing protein production ^4^. As master regulators of gene expression, miRNA have been investigated in multiple pathologies to identify their functional pathways and biomarker potential ^5,6^. During mycobacterial infections, miRNA are differentially regulated, and play significant biological roles during infection ^7–11^. The regulation of miRNA expression is a contested node, in that it is multifaceted and driven by several opposing factors. Host control of miRNA is challenged by pathogen-driven modulation of expression and the downstream impacts on host immunity may result in either successful clearance or subversion by the pathogen to support infection.

Reduced expression of miR-126 has been reported in cattle suffering from Johne’s disease, an infection caused by *Mycobacterium avium* subspecies *paratuberculosis*, in *Mycobacterium abscessus* and *Mycobacterium tuberculosis*-infected THP-1 cells, and in patients with tuberculous meningitis, a severe manifestation of *M*. *tuberculosis* infection, and pulmonary tuberculosis. ^12–16^. Although mapped targets suggest further roles in haematopoiesis and inflammatory disorders, the function of miR-126 in infection has yet to be defined ^17–20^.

Zebrafish (*Danio rerio*) are a powerful model organism for studying host-mycobacterial interactions, as they allow direct visualisation of cellular interactions and are amenable to straightforward genetic manipulation ^1^. An added benefit of the zebrafish-*M*. *marinum* model is the ability to use a native pathogen of the zebrafish, which closely mirrors histological aspects of human-*M*. *tuberculosis* pathogenesis ^21,22^. Zebrafish provide an established model for the investigation of miRNA function and responses to infection, with well-developed genetic tools that facilitate the study of downstream host gene function. We have recently used the zebrafish-*M*. *marinum* model to investigate the downstream targets of miR-206 and identified the role of this conserved miR in controlling the multi-cellular immune response to mycobacterial infection ^7^.

Here we use the zebrafish-*M*. *marinum* platform to examine the role of mycobacterium infection-induced dre-miR-126a-3p (miR-126). We link infection-induced miR-126 to Tsc1 via activation of the mTOR pathway, and improved macrophage function due to regulation of a Cxcl12/Ccl2/Ccr2 signalling axis during the early stages of mycobacterial infection.

## Results

### Infection-induced miR-126 is host-protective against *M*. *marinum* infection

Embryos were infected with *M*. *marinum* via caudal vein injection and analysed at 1- and 3-days post infection (dpi) by quantitative (q)PCR to assess the responsiveness of miR-126 expression to infection. At both timepoints miR-126 was upregulated in *M*. *marinum*-infected embryos in comparison to uninfected control embryos (Figure 1A).

**Figure 1.**
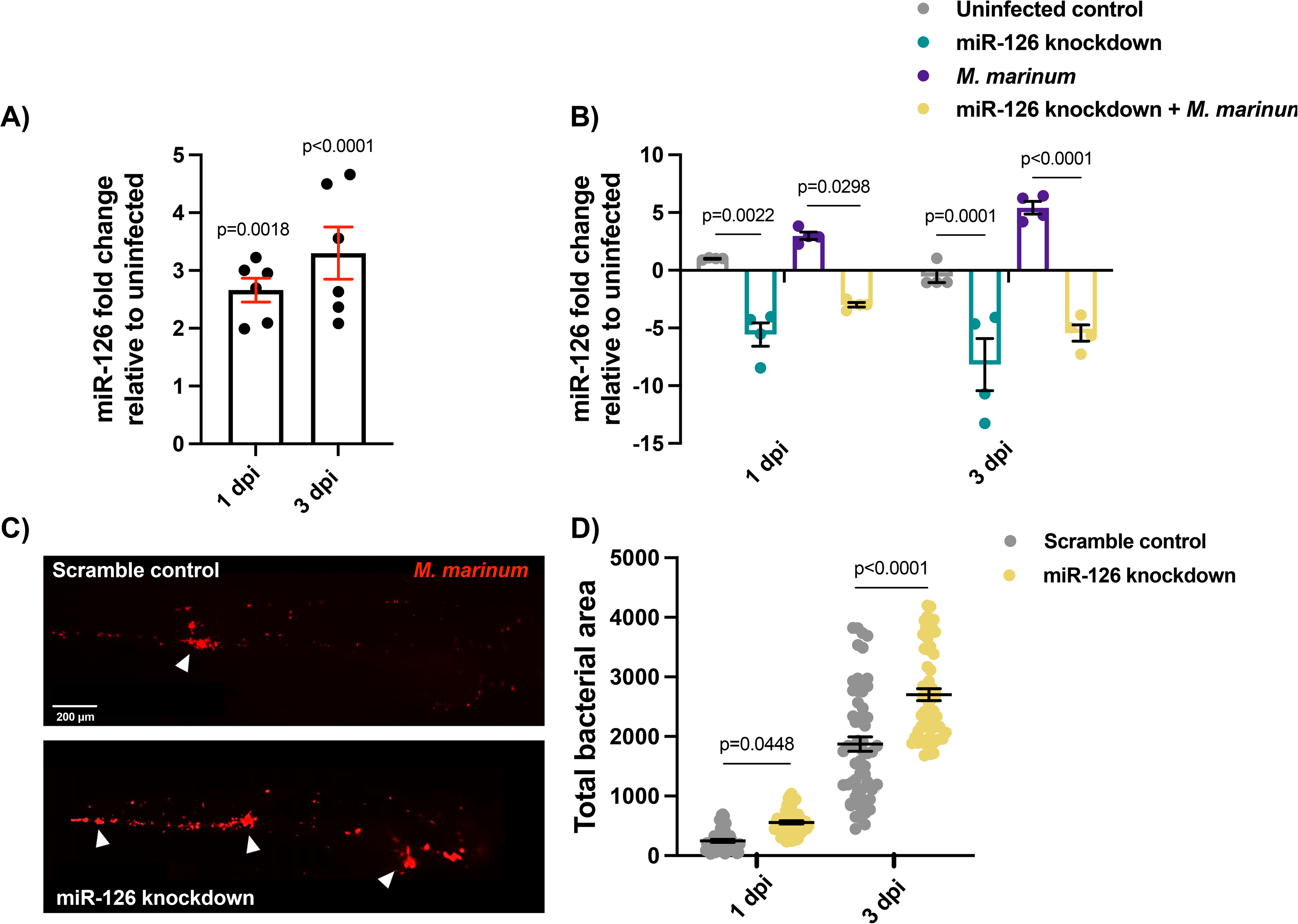
Infection-induced miR-206 expression alters bacterial burden. (A) Expression of miR-206 following *M*. *marinum* infection analysed by qPCR at 1 and 3 dpi relative to uninfected embryos. (B) Expression of miR-206 in uninfected and infected, antagomir-injected embryos (miR-206 knockdown). (C) Representative images of *M*. *marinum* infection at 3 dpi in control and miR-126 knockdown embryos. Scale bar represents 200 μm. (D) *M*. *marinum* burden in miR-206 knockdown embryos at 1 and 3 dpi. Each data point represents a single measurement, with the mean and SEM shown. For qPCR analysis, each data point represents 10 embryos, and contains 2 biological replicates. Bacterial burden analysis data points represent individual embryos (n=40-50 embryos per group) and are representative of 2 biological replicates.

To determine if antagomiR-mediated knockdown of miR-126 effectively reduced transcript abundance and if this miR-126 knockdown was sustained following *M*. *marinum* infection, embryos were injected with antagomiR at the single-cell stage and infected with *M*. *marinum* at 1.5 days post fertilisation (dpf). Expression levels of miR-126 were measured at 1 and 3 dpi. AntagomiR knockdown reduced miR-126 levels at both timepoints and prevented the infection-associated miR-126 expression (Figure 1B).

The effect of miR-126 expression on mycobacterial infection was assessed through analysis of bacterial burden following infection in control and miR-126 knockdown embryos (Figure 1C). There was a small, but statistically significant, increase in the bacterial burden of miR-126 knockdown embryos at 1 dpi, which was much more apparent at 3 dpi (Figure 1D). These results indicate that *M*. *marinum* infection induces upregulation of miR-126 and that miR-126 has a host-protective effect.

### Infection-associated miR-126 expression is enhanced by mycobacterial virulence factors

To investigate whether the increased bacterial burden observed in miR-126 knockdown embryos was specific to mycobacterial infection or a universal response to invading pathogens, embryos were infected with ΔESX1 *M*. *marinum* or uropathogenic *Escherichia coli* (UPEC). ΔESX1 *M*. *marinum* lack a key mycobacterial type VII secretion system and are therefore less virulent than wild-type (WT) *M*. *marinum* as they are unable to escape host phagosomes (Conrad et al. 2017). UPEC are a predominately extracellular bacterium that, in contrast to intracellular mycobacteria, cause acute sepsis infection ^23^.

Expression of miR-126 was analysed by qPCR at 1 dpi following infection with either *M*. *marinum*, ΔESX1 *M*. *marinum*, or UPEC. While infection with either ΔESX1 *M*. *marinum* or UPEC increased miR-126 expression in comparison to control uninfected embryos, the level of miR-126 expression was significantly lower than WT virulent *M*. *marinum* (Figure 2A).

**Figure 2.**
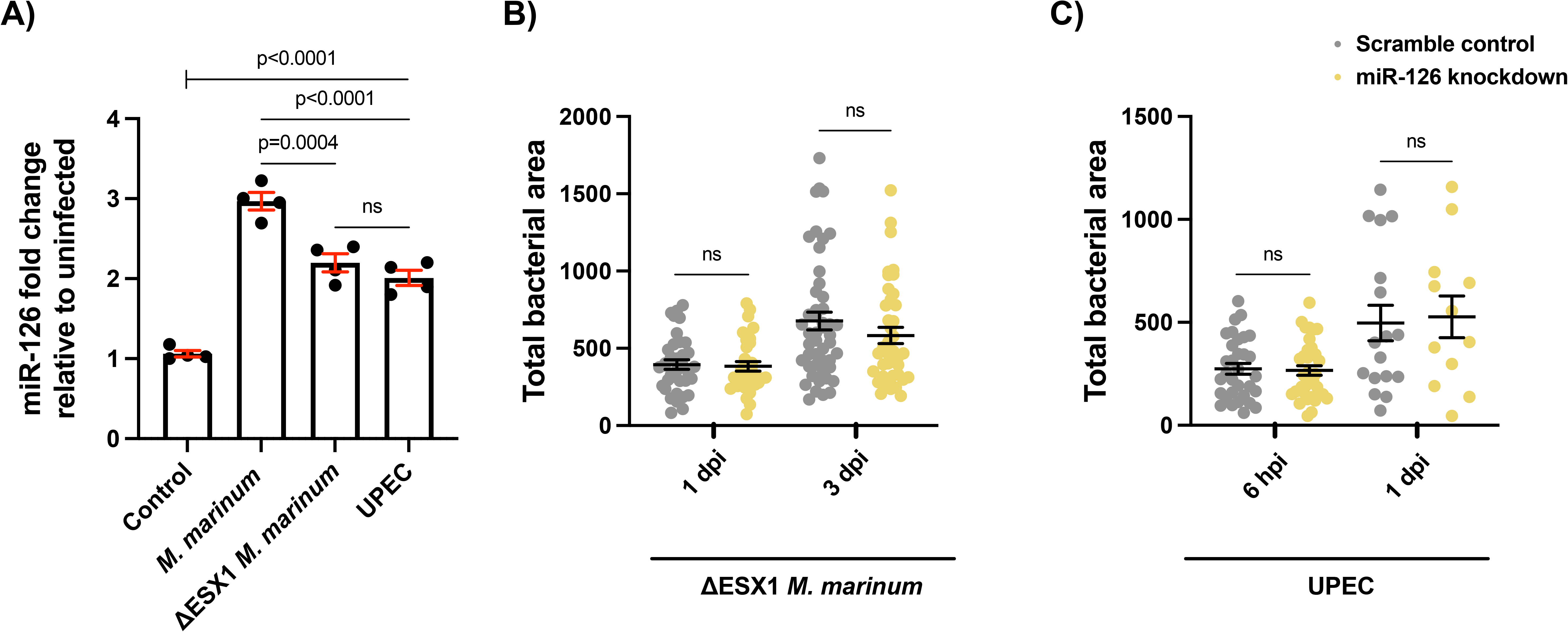
Mycobacterial virulence factors drive the induction and protective effects of miR-126. (A) Expression of miR-126 at 1 dpi following infection with either WT *M*. *marinum*, ΔESX1 *M*. *marinum*, or UPEC. (B) ΔESX1 *M*. *marinum* burden at 1 and 3 dpi in miR-126 knockdown embryos. (C) UPEC burden at 6 hpi and 1 dpi in miR-126 knockdown embryos. For qPCR analysis, data points are representative of a single measurement of 10 pooled embryos and 2 experimental replicates, with the mean and standard error of the mean (SEM) shown. For bacterial burden analysis, each data point represents a single measurement (n=35-45 embryos per group for ΔESX1 *M*. *marinum* and n=15-35 embryos per group for UPEC), and 2 experimental replicates with the mean and SEM shown.

Knockdown of miR-126 did not affect ΔESX1 *M*. *marinum* burden compared to control embryos at 1 or 3 dpi (Figure 2B). Likewise, there was no difference in UPEC burden between control and miR-126 knockdown embryos at either 6 hours post infection (hpi) or 1 dpi (Figure 2C). These results suggest that although the induction of miR-126 is conserved between types of infections, the host-protective action of miR-126 is only uncovered in the context of infection with virulent *M*. *marinum*.

### miR-126 target gene mRNA expression patterns are conserved during *M*. *marinum* infection of zebrafish

To uncover the biological pathways targeted by miR-126, expression of potential target mRNAs was analysed by qPCR following antagomiR knockdown and infection with *M*. *marinum*. Possible target genes were chosen based on published experimentally observed targets and bioinformatic target prediction software (Supplementary File 1) ^24–34^. Increased expression of a transcript in miR-126 knockdown embryos compared to uninfected scrambled controls was expected to indicate targeting by miR-126 (Figure 3A-E).

**Figure 3.**
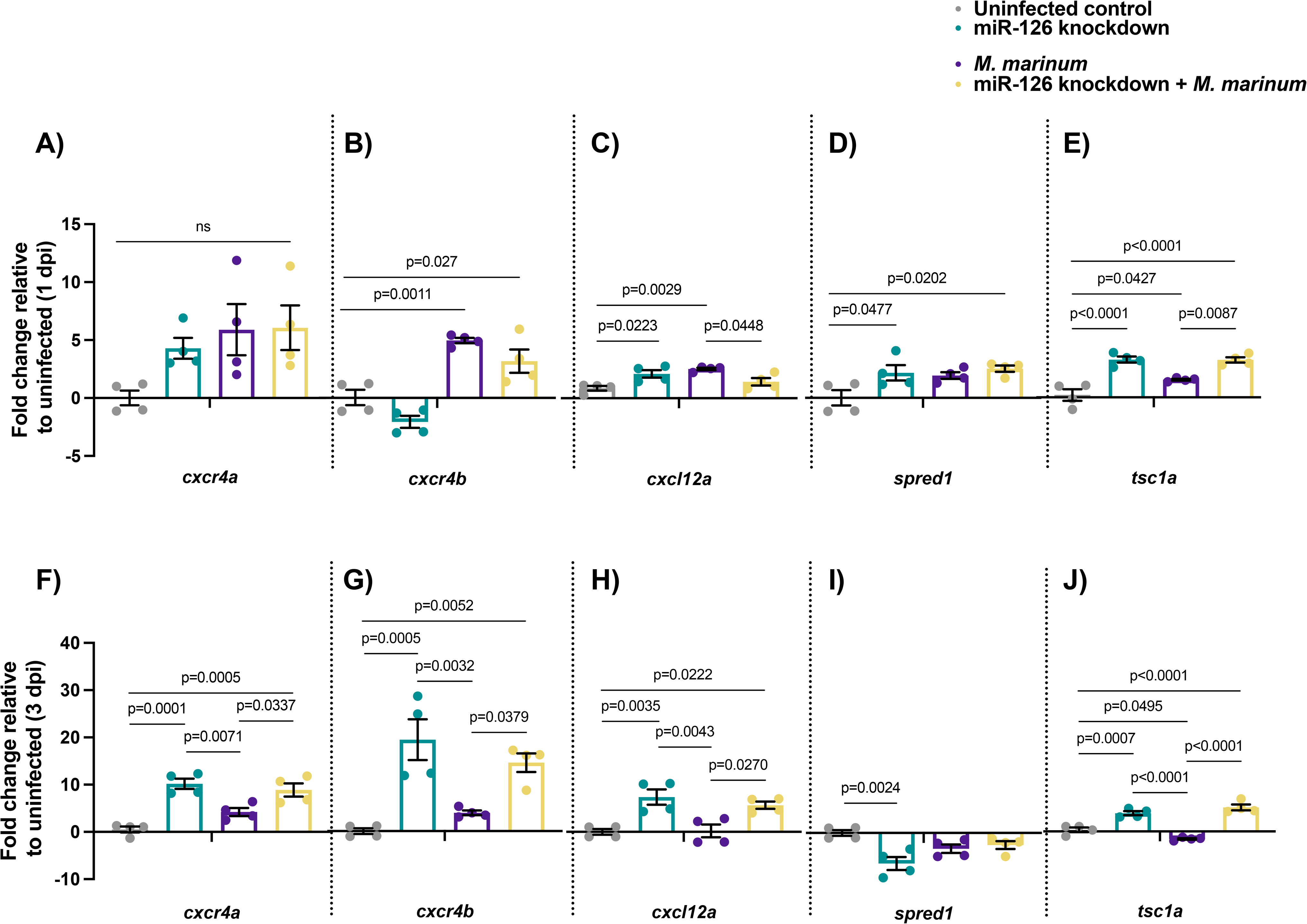
Expression of potential miR-126 mRNA targets is conserved in zebrafish *M*. *marinum* infection. (A-E) Expression of candidate genes at 1 dpi in miR-126 knockdown measured by qPCR. (F-J) qPCR analysis of zebrafish CXR4/CXCL12 and TSC1 ortholog genes at 3 dpi. Each data point represents a single measurement of 10 pooled embryos and 2 technical replicates, with the mean and SEM shown.

Expression of *cxcr4b* was significantly increased in *M*. *marinum* infected and miR-126 knockdown + *M*. *marinum* embryos compared to control at 1 dpi. Expression of *cxcr4a* was increased in all treatments but was not significantly different from control uninfected embryos. Expression of *cxcl12a* was higher in both miR-126 knockdown and *M*. *marinum* infected embryos than control, but not knockdown infected embryos. miR-126 knockdown + *M*. *marinum* embryos had reduced *cxcl12a* expression compared to control infected embryos. While expression of *spred1* was increased in both knockdown groups compared to control, there was no difference between knockdown infected and control infected embryos. Finally, *tsc1a* was increased in all treatments from control embryos, while expression was significantly higher in miR-12 knockdown and miR-126 knockdown + *M*. *marinum* embryos than *M*. *marinum* alone.

From the gene expression analysis, *cxcr4b*, *cxcl12a*, and *tsc1a* were regarded as potential target genes of miR-126 due to their increased expression in knockdown embryos and differential regulation compared to *M*. *marinum* infected embryos. Both *cxcr4a/b* and *cxcl12a* were of interest due to their previously identified role in mycobacterial and zebrafish immunity ^7,35,36^. Also of interest, *tsc1a* may be involved in mycobacterial pathogenesis as a negative regulator of the mTOR signalling pathway, a key regulatory pathway of a variety of cellular functions ^37,38^. Expression of *cxcr4a, cxcr4b, cxcl12a*, and *tsc1a* was measured at 3 dpi, and showed that knockdown of miR-126 increased the expression of all target genes even in *M*. *marinum*-infected knockdown embryos, when compared to control uninfected and *M*. *marinum*-infected only embryos (Figure 3F-J).

### miR-126 suppression of tsc1a expression aids control of *M. marinum* infection

Following expression profiling, *tsc1a* was selected for further investigation due to differential expression between *M*. *marinum* infected and miR-126 knockdown embryos. Predicted binding of miR-126 and *tsc1a* in zebrafish is summarised in (Figure 4A). To confirm the interaction between miR-126 and *tsc1a*, we targeted *tsc1a* for knockdown using CRISPR-Cas9 (Figure 4B). Knockdown of *tsc1a* significantly reduced transcript abundance at 1 and 3 dpi compared to both control uninfected and *M*. *marinum*-infected embryos (Figure 4C). At both timepoints, *tsc1a* knockdown was sustained in *M*. *marinum*-infected knockdown embryos. Knockdown of *tsc1a* significantly reduced *M*. *marinum* burden compared to control embryos (Figure 4D). Double knockdown of both miR-126 and *tsc1a* significantly reduced bacterial burden compared to miR-126 knockdown alone. There was no difference in bacterial burden between double knockdown embryos and *tsc1a* knockdown alone embryos, suggesting that *tsc1a* is driving the miR-126 knockdown-associated increase in burden (Figure 4D). The opposing effects of miR-126 knockdown and *tsc1a* knockdown on *M*. *marinum* burden suggested involvement of a potential miR-126/Tsc1 signalling axis in mycobacterial infection.

**Figure 4.**
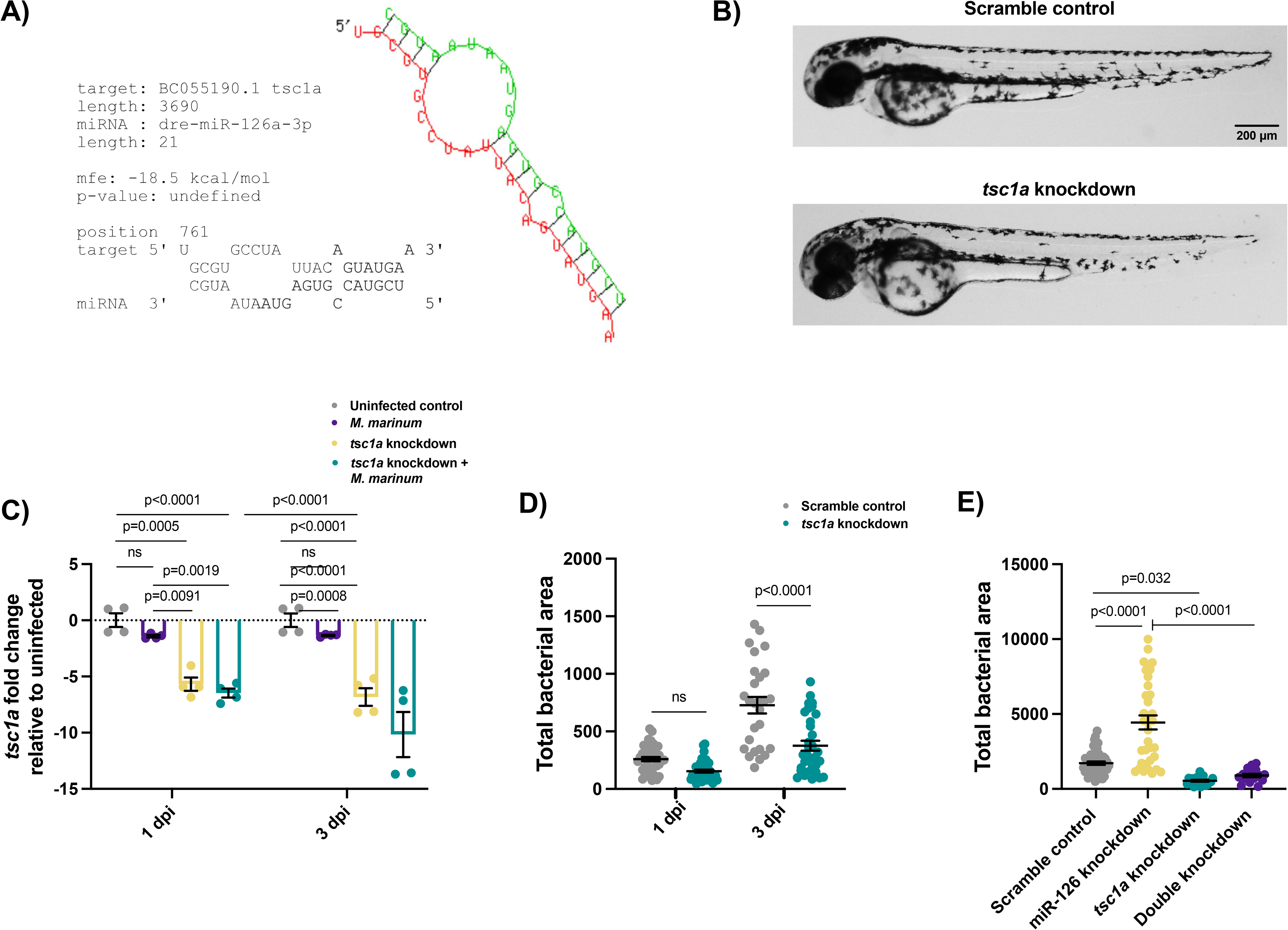
miR-126 targets *tsc1a* to worsen *M*. *marinum* infection burden. (A) Binding kinetics of *tsc1a* and dre-miR-126a-3p as predicted by RNAhybrid. (B) Brightfield images of *tsc1a* knockdown embryos at 3 dpf showing no abnormal developmental phenotypes. Scale bar represents 200 μm. (C) *tsc1a* expression was measured by qPCR at 1 and 3 dpi following CRISPR-Cas9 knockdown of *tsc1a* and infection with *M*. *marinum*. (D) *tsc1a* knockdown embryos were infected with *M*. *marinum* via caudal vein injection and bacteria burden was analysed at 1 and 3 dpi. (E) *tsc1a* and miR-126 double knockdown embryos were infected with *M*. *marinum* via caudal vein injection and bacterial burden was analysed at 3 dpi. For qPCR analysis, data points are representative of a single measurement of 10 pooled embryos and 2 experimental replicates, with the mean and standard error of the mean (SEM) shown. Bacterial burden data points represent a single measurement (n=27-44 embryos per group [C] and n=11-20 embryos per group [D]), and 2 experimental replicates with the mean and SEM shown.

### miR-126 knockdown alters the mTOR signalling axis in *M*. *marinum* infection

As miR-126 knockdown increased expression of *tsc1a*, which encodes a negative regulator of mTOR function, we investigated downstream mTOR activity. We utilised whole-mount embryo immunofluorescent staining of phosphorylated ribosomal protein S6 (phospho-S6) as a readout for mTOR activity following infection of miR-126 and *tsc1a* knockdown embryos with *M*. *marinum* (Figure 5A). At 1 dpi, knockdown of miR-126 resulted in reduced phospho-S6 staining, consistent with increased *tsc1a* transcript abundance and knockdown of either *tsc1a* alone or double knockdown of miR-126 and *tsc1a* increased phospho-S6 staining compared to both control and miR-126 knockdown embryos (Figure 5B). However, the bulk changes in phosphor-S6 staining occurred largely distal to the fluorescent *M*. *marinum*, with no observed colocalization, suggesting a lack of specificity to infection (Figure 5A).

**Figure 5.**
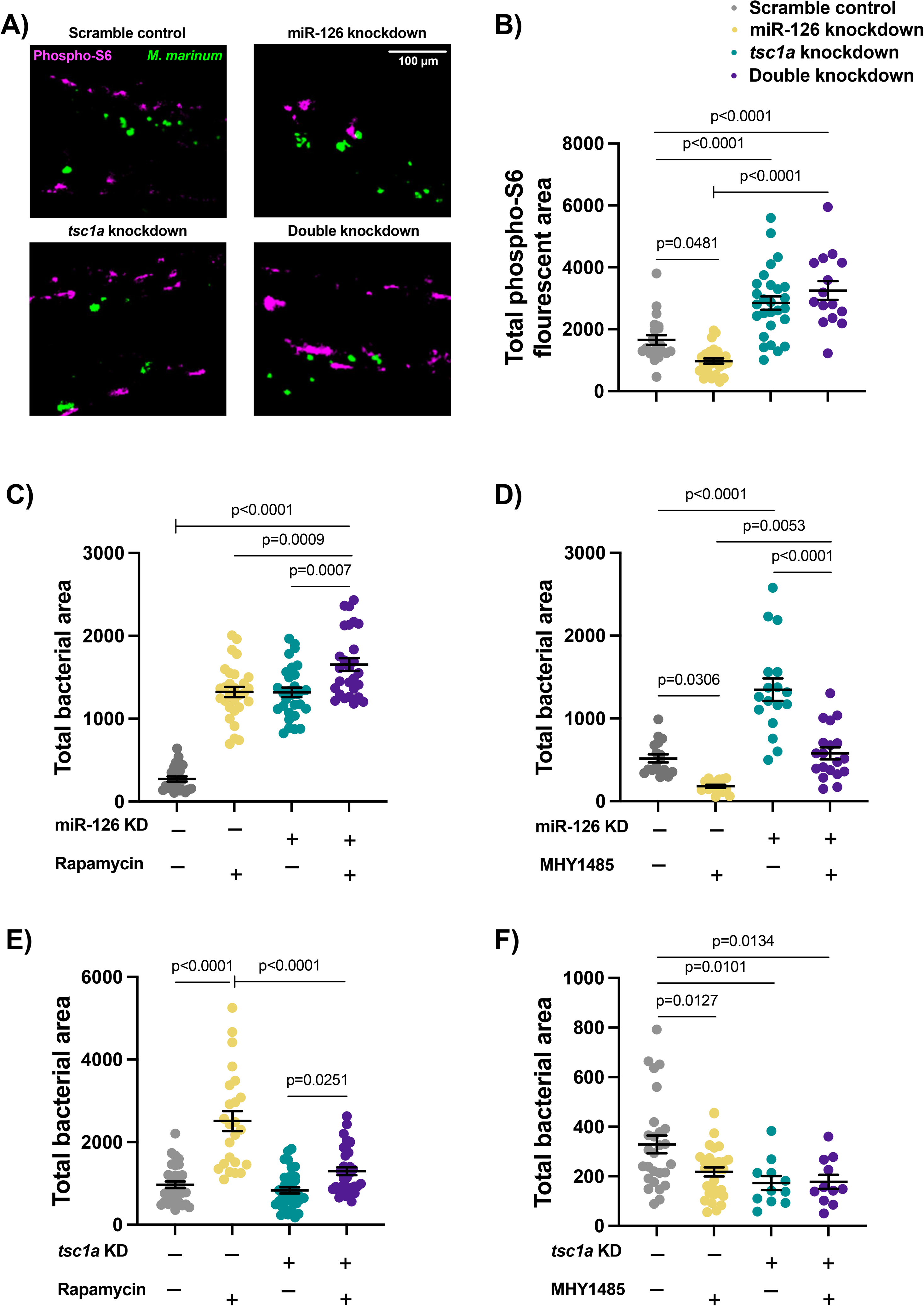
miR-126 acts on *tsc1a* to influence mTOR signalling during infection. (A) Representative images of phospho-S6 fluorescent staining in miR-126 and *tsc1a* knockdown embryos at 1 dpi. *M*. *marinum* is green and phosphorylated ribosomal protein S6 is magenta. Scale bar represents 100 m. (B) Phospho-S6 staining in *M*. *marinum*-infected control, miR-126 knockdown, and *tsc1a* knockdown at 1 dpi (C) miR-126 knockdown embryos were infected with *M*. *marinum* via caudal vein injection and treated with mTOR inhibitor rapamycin. Bacterial burden was analysed at 1 dpi. (D) miR-126 knockdown embryos were infected with *M*. *marinum* via caudal vein injection and treated with mTOR activator MHY1485. (E) *tsc1a* knockdown embryos were infected with *M*. *marinum* via caudal vein injection and treated with mTOR inhibitor rapamycin. Bacterial burden was analysed at 1 dpi. (F) *tsc1a* knockdown embryos were infected with *M*. *marinum* via caudal vein injection and treated with mTOR activator MHY1485. Bacterial burden was analysed at 1 dpi. Bacterial burden was analysed at 1 dpi. Each data point represents a single measurement (n=11-36 embryos per group) with the mean and SEM shown. Phospho-S6 staining is a single experimental replicate, while rapamycin and MHY1485 treatments represent 2 experimental replicates each.

To assess the impact of decreased mTOR signalling in miR-126 knockdown, embryos were treated with either an inhibitor (rapamycin) or activator (MHY1485) of mTOR. Treatment of control embryos with rapamycin resulted in a similar increased burden to that seen after knockdown of miR-126 (Figure 5C). The combination of miR-126 knockdown and rapamycin treatment had an additive effect on the increased bacterial burden compared to either treatment alone, suggesting the presence of infection-relevant, non-mTOR miR-126 activities.

Conversely, activation of mTOR signalling with MHY1485 significantly reduced the bacterial burden in both control and miR-126 knockdown backgrounds (Figure 5D). The combination of miR-126 knockdown and MHY1485 treatment only partially rescued the miR-126 knockdown-induced increase in bacterial burden as compared to MHY1485 treatment alone, again indicating that these pathways are not fully interlinked and that multiple miR-126 dependant mechanisms are functioning during infection.

As anticipated, inhibition of mTOR signalling through rapamycin treatment of *tsc1a* knockdown embryos increased bacterial burden, however not to the levels seen in rapamycin-treated scramble control embryos (Figure 5E). Also as anticipated, treatment with MHY1485 did not further decrease burden levels from *tsc1a* knockdown alone (Figure 5F). Together, these data potentially place mTOR activation as one of the factors downstream of infection-induced miR-126 repression of *tsc1a* expression and demonstrate that additional non-mTOR factors mediate the infection phenotype downstream of the miR-126/*tsc1a* axis.

### miR-126 knockdown and inhibition of mTOR increase cell death in *M*. *marinum* infection

Because of the pleiotropic effects of mTOR on apoptosis and cell death, we hypothesised that increased mTOR activity caused by increased *tsc1a* expression in miR-126 knockdown embryos might compromise the survival of *M*. *marinum*-infected macrophages ^39–41^. To explore this hypothesis, we performed TUNEL staining on 3 dpi miR-126 knockdown embryos (Figure 6A). Knockdown of miR-126 did increase the number of TUNEL-stained cells compared to control infected embryos (Figure 6B).

**Figure 6.**
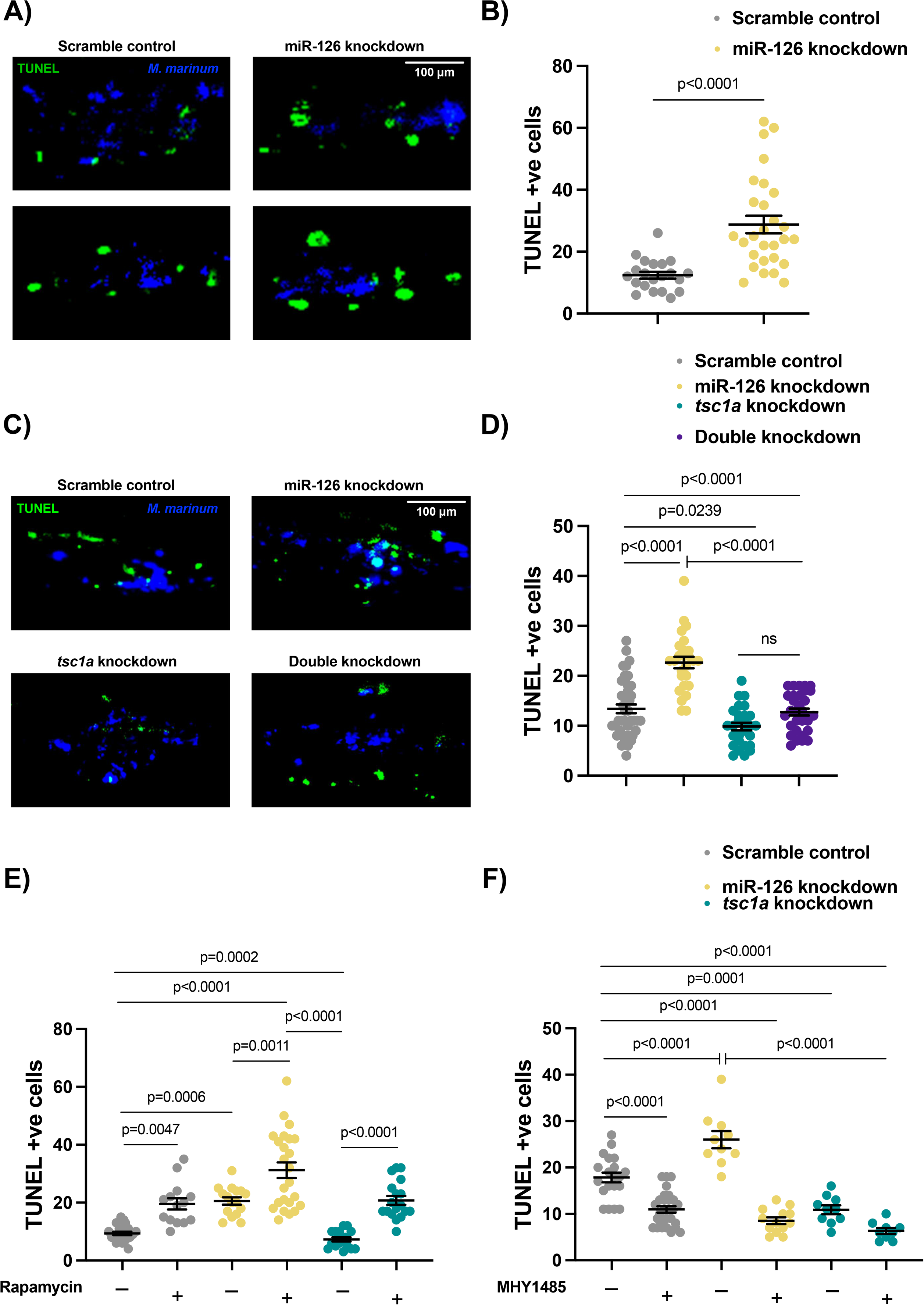
Decreased mTOR signalling alters cell death dynamics in mycobacterial infection. (A) Representative images of TUNEL staining in miR-126 knockdown embryos at 3 dpi. TUNEL +ve cells are green and *M*. *marinum* is blue. Scale bar represents 100 μm. (B) TUNEL +ve cells in *M. marinum*-infected control and miR-126 knockdown embryos were counted at 3 dpi. (C) Representative images of TUNEL staining in *tsc1a* and double knockdown embryos at 3 dpi. TUNEL +ve cells are green and *M*. *marinum* is blue. Scale bar represents 100 μm. (D) TUNEL +ve cells in *M*. *marinum*-infected *tsc1a* and double knockdown embryos were counted at 3 dpi. (E) TUNEL +ve cells in rapamycin-treated, *M*. *marinum*-infected knockdown embryos were counted at 3 dpi. (F) TUNEL +ve cells in MHY1485-treated, *M*. *marinum*-infected knockdown embryos were counted at 3 dpi. Each data point represents a single measurement, with the mean and SEM shown (n=20-25 embryos per group). Graphs are representative of 2 experimental replicates with the exception of rapamycin/MHY1485 experiments, which were performed in a single replicate.

The increased cell death in miR-126 knockdown embryos was completely reversed by double knockdown of miR-126 and *tsc1a* indicating the cell death phenotype is dependent on *tsc1* expression (Figure 6C-D). Rapamycin treatment increased the number of TUNEL-stained cells in control and *tsc1a* knockdown embryos, and in the miR-126 knockdown embryos as expected from the bacterial burden data (Figure 6E).

Although these data confirm the independence between miR-126/*tsc1a* and mTOR activity during mycobacterial infection, the increased cell death in miR-126 knockdown embryos was completely suppressed by treatment of miR-126 knockdown embryos with the mTOR activator MHY1485 demonstrating a role for mTOR activity in regulating cell death at the host-mycobacterial interface (Figure 6F).

### miR-126 knockdown increases the migration of macrophages to sites of *M*. *marinum* infection

As knockdown of miR-126 also increased expression of the neutrophil-related genes, *cxcr4b* and *cxcl12a*, and we had previously found a role for upregulation of these genes in protection from *M*. *marinum* infection ^7^, we investigated the effect of miR-126 knockdown on neutrophil motility. Neutrophil responses were first assessed via static imaging of whole-body neutrophil numbers and then by time-lapse imaging of trunk-infected embryos for analysis of neutrophil migration in the transgenic (Tg) *Tg(lyzC:GFP)^nz117^* neutrophil reporter line. Static imaging revealed no difference in total neutrophil numbers between *M*. *marinum*-infected control and miR-126 knockdown embryos at either timepoint (Figure 7A). Migration of neutrophils to the site of infection was also not altered between miR-126 knockdown and control infected embryos (Figure 7B). This suggests that despite the increased abundance of transcripts encoding the neutrophil chemotactic genes *cxcr4a/b* and *cxcl12a* in miR-126 knockdown embryos, neutrophil recruitment is unperturbed by miR-126 knockdown in *M*. *marinum*-infected embryos.

**Figure 7.**
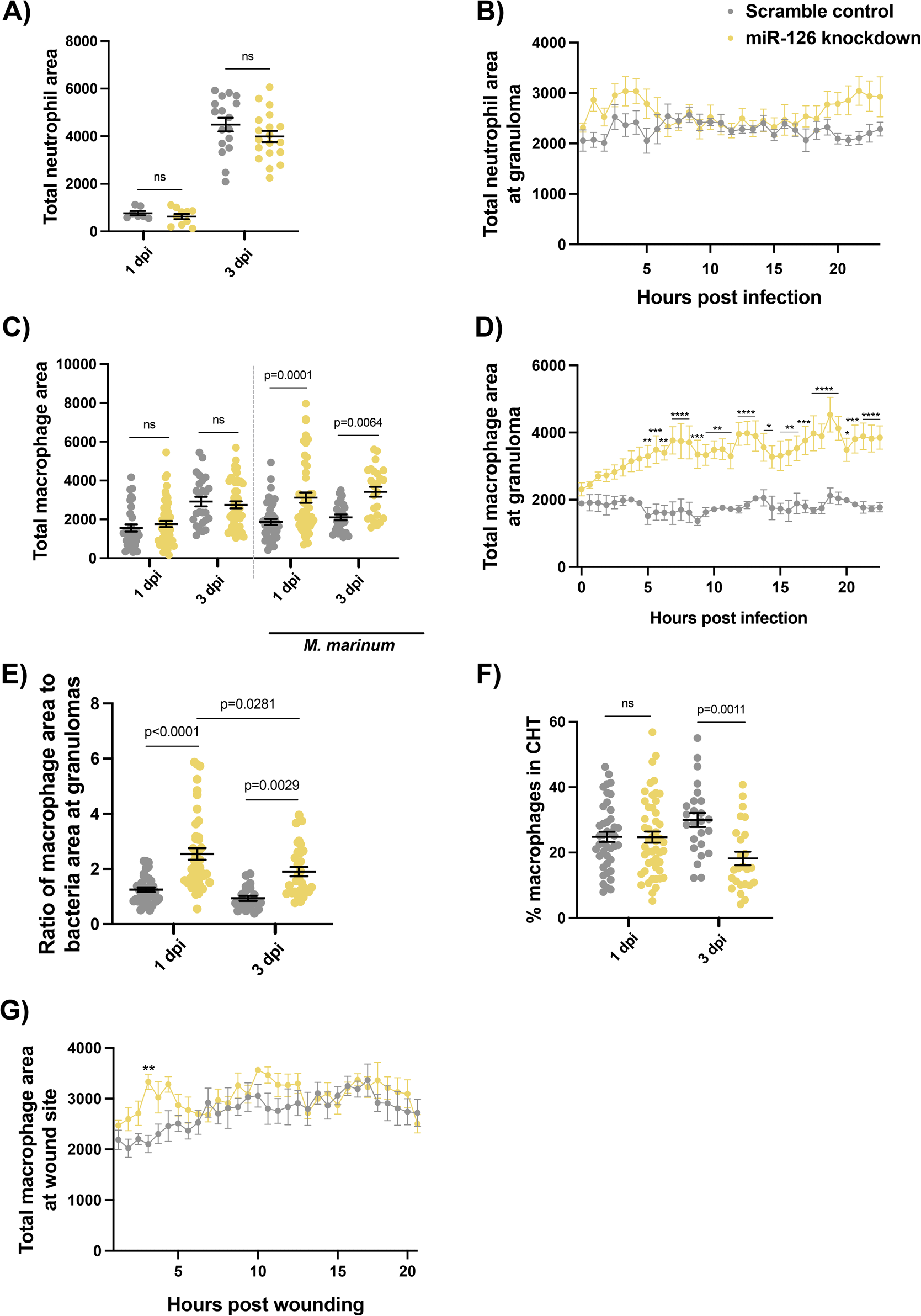
Mycobacterial infection-induced miR-126 expression alters the host macrophage response. (A) Measurement of whole-body neutrophil fluorescent area at 1 and 3 dpi in control and miR-126 knockdown infected embryos. (B) Measurement of neutrophil levels following trunk infection with *M*. *marinum* in miR-126 knockdown embryos. (C) Measurement of whole-body macrophage fluorescent area at 1 and 3 dpi in uninfected and infected miR-126 knockdown embryos. (D) Measurement of macrophage levels following trunk infection with *M*. *marinum* in miR-126 knockdown embryos. (E) Ratio of macrophage fluorescent area per bacterial fluorescent area at granulomas in miR-126 knockdown embryos at 1 and 3 dpi. (F) Measurement of macrophage recruitment to a tail wound in miR-126 knockdown embryos. Each data point represents a single measurement with the mean and SEM shown. For neutrophil analysis 10-20 embryos per group were analysed, and 15-50 embryos per group for macrophage analysis. For neutrophil time-lapse imaging, each data point represents the mean of 6 foci of infection from 6 separate embryos, and the graph is representative of 2 experimental replicates. For macrophage time-lapse imaging, each data point represents the mean of 3 foci of infection from 3 separate embryos, and the graph is representative of 2 experimental replicates. * P < 0.05, ** p < 0.01, *** p < 0.001, **** p < 0.0001.

Because of the role of CXCL12 in recruiting CCR2-expressing macrophages and directing their function towards anti-inflammatory activities ^42,43^, we next investigated the macrophage response to *M*. *marinum* infection in miR-126 knockdown embryos. We first estimated total macrophage number in *Tg(mfap4:turquoise)^xt27^* and *Tg(mfap4:tdtomato)^xt12^* macrophage reporter lines embryos at baseline and following *M*. *marinum* infection. While there was no difference in total macrophage number between control and miR-126 knockdown embryos in the absence of infection, *M*. *marinum*-infected miR-126 knockdown embryos had significantly more macrophages than control infected embryos (Figure 7C).

To track the recruitment of macrophages to discrete *M*. *marinum* lesions, embryos were infected with *M*. *marinum* via a trunk injection that embeds the bacteria away from the caudal haematopoietic tissue (CHT) (Supplementary videos 1-2). Compared to control embryos, miR-126 knockdown embryos had an increased number of macrophages at the site of infection from 5 to 24 hpi (Figure 7D). The increased association of macrophages with *M*. *marinum* was maintained at 3 dpi in miR-126 knockdown embryos (Figure 7E). There was initially no difference in the number of macrophages present in the CHT at 1 dpi, however CHT macrophage numbers were decreased in miR-126 knockdown embryos at 3 dpi, suggesting that reduced miR-126 increases the migration of macrophages but does not influence the production and maturation of immature progenitors (Figure 7F).

In order to determine if the increased migratory potential of macrophages in miR-126 knockdown embryos was specific to infection, we performed a sterile tail fin wounding assay (Supplementary videos 3-4). There was no significant difference in the number of macrophages at the wound site between miR-126 knockdown and control embryos, suggesting the alteration of the macrophage response dynamics is dependent on specific pathogen-derived signals (Figure 7G).

### Increased macrophage recruitment to *M*. *marinum* infection is independent of Tsc1a induction in miR-126 knockdown embryos

We next sought to determine if the Tsc1a/mTOR axis affected macrophage recruitment. Following infection with *M*. *marinum, tsc1a* knockdown embryos did not display any difference in total macrophage numbers compared to control embryos at either 1 or 3 dpi (Fig 8A). Double knockdown of miR-126 and *tsc1a* failed to prevent the increased macrophage numbers seen in miR-126 knockdown at 1 dpi, indicating that the altered macrophage response is independent of Tsc1a/mTOR (Fig 8B).

**Figure 8.**
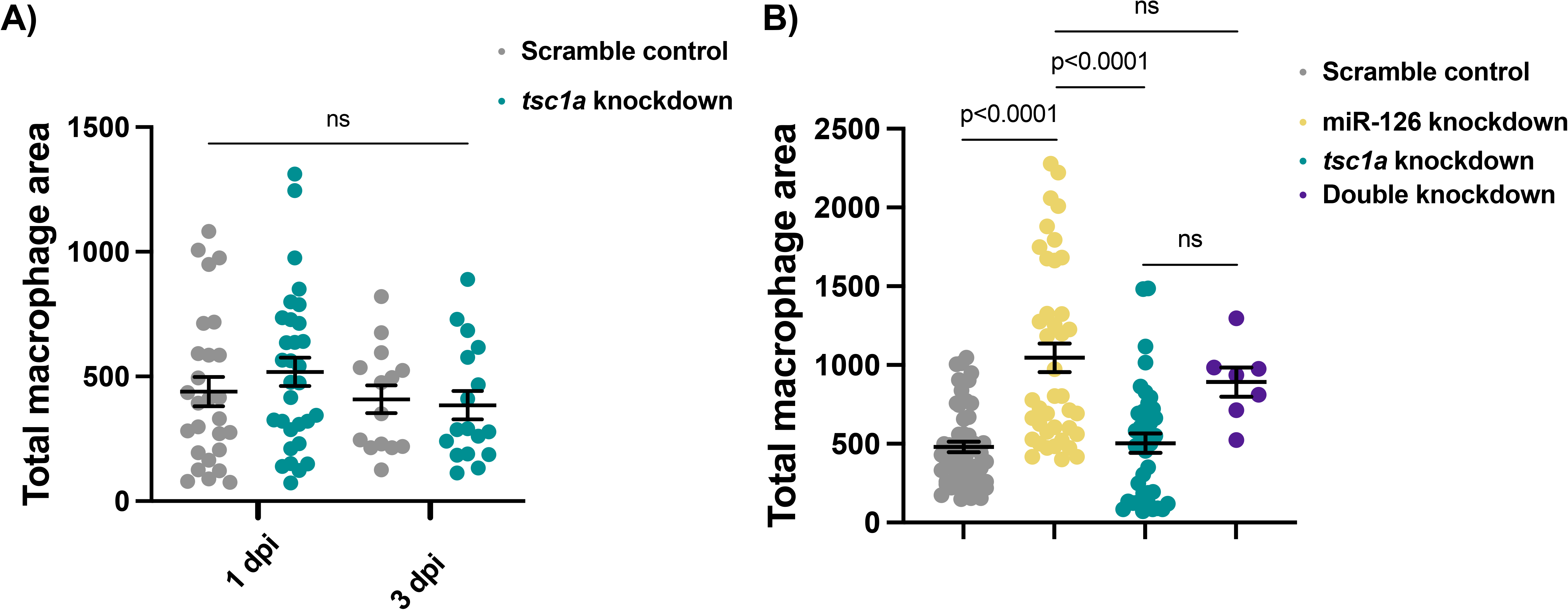
miR-126-dependent macrophage responses to infection are not controlled by the Tsc1a/mTOR signalling axis. (A) Measurement of whole-body macrophage fluorescent area at 1 and 3 dpi in *M*. *marinum*-infected control and tsc1a knockdown embryos. (B) Measurement of whole-body macrophage fluorescent area at 1 dpi in *M*. *marinum*-infected control and knockdown embryos. Each data point represents a single measurement, with the mean and SEM shown (n=7-56) embryos per group). Graph is representative of 2 experimental replicates.

### Recruited macrophages are permissive to *M*. *marinum* infection

To determine why miR-126 knockdown embryos were unable to contain *M*. *marinum* in granulomas, despite increased macrophage recruitment, we assessed the activation state of the macrophages. We hypothesised that increased *cxcl12* expression provides more ligand, by heterodimerization with Ccl2, also known as monocyte chemoattractant protein-1 (Mcp-1), for the recruitment of Ccr2 positive (Ccr2+) permissive or non-bactericidal macrophages to sites of infection ^42^.

To investigate this, we first assessed *ccl2* and *ccr2* transcript abundance following infection. Knockdown of miR-126 increased expression of *ccr2* compared to scramble control embryos (Figure 9A). Expression of *ccr2* was also increased when miR-126 knockdown embryos were infected with *M*. *marinum* compared to infected, scramble control embryos. While *ccl2* expression was not responsive to either miR-126 knockdown or infection alone, infection of miR-126 knockdown embryos increased *ccl2* expression compared to knockdown alone and *M. marinum*-infected only embryos (Figure 9B). Further, *ccl2* but not *ccr2* expression was observed to be dependent on *cxcl12a* expression, with double knockdown of miR-126 and *cxcl12a* rescuing *ccl2* expression (Supplementary Figure 1).

**Figure 9.**
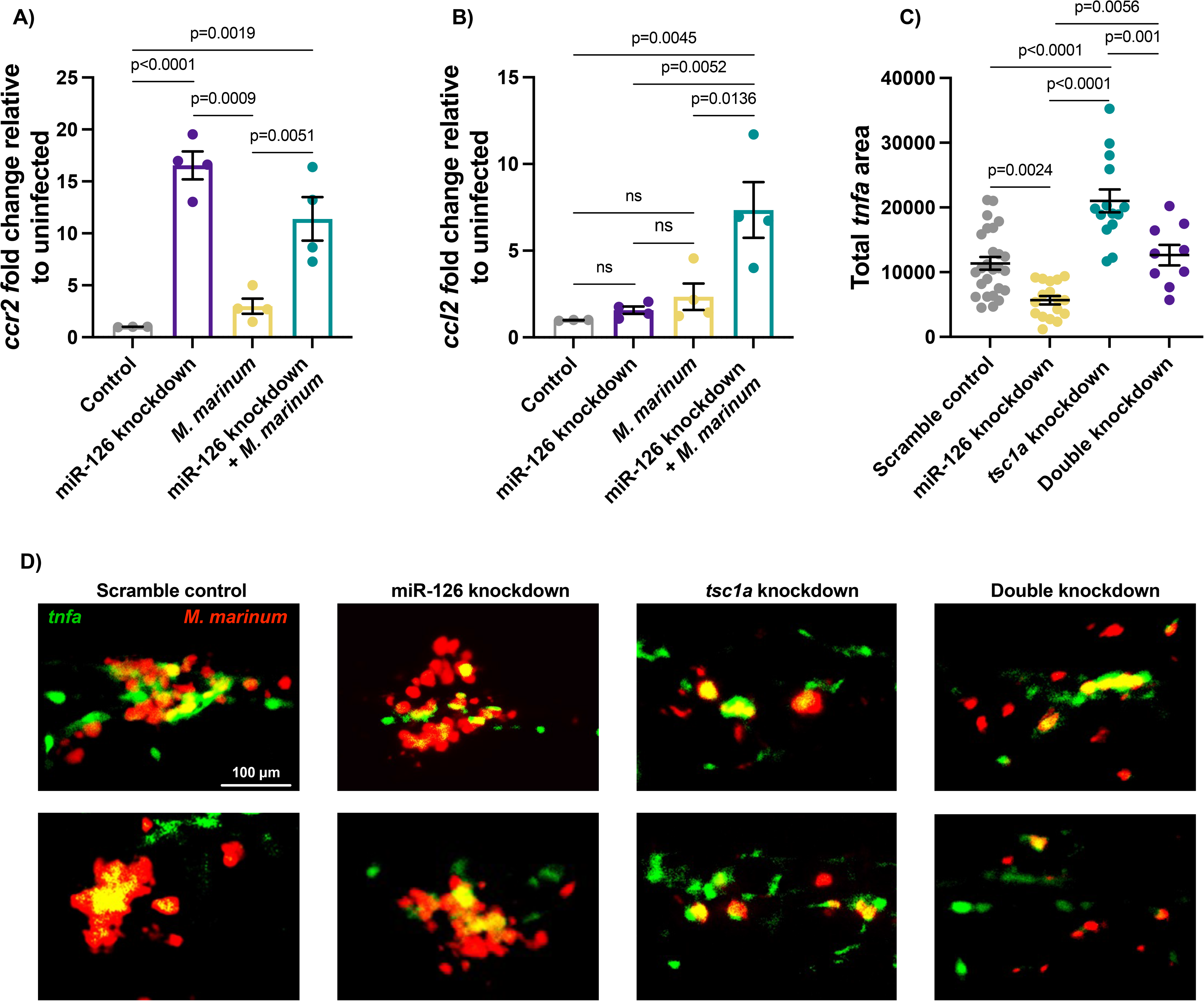
Mycobacterial infection-induced miR-126 expression increases proinflammatory bactericidal macrophage recruitment. (A-B) Expression of *ccr2* and *ccl2* was analysed by qPCR at 1 dpi following miR-126 knockdown and infection with *M*. *marinum* (C) Measurement of *tnfa* promoter activation following trunk infection with *M*. *marinum* in knockdown embryos at 1 dpi. infected with *M*. *marinum* via caudal vein injection and bacterial burden was analysed at 3 dpi (D) Representative images of *tnfa* promoter-drive GFP expression at 1 dpi in knockdown embryos following trunk infection with *M*. *marinum*. *M*. *marinum* is red, and *tnfa* is green. Scale bar represents 100 m. For qPCR analysis, data points are representative of a single measurement of 10 pooled embryos and 2 experimental replicates, with the mean and standard error of the mean (SEM) shown. For bacterial burden, each data point represents a single measurement, with the mean and SEM shown (n=9-26) embryos per group). Graphs are representative of 2 experimental replicates.

To determine if recruited macrophages were alternatively activated, and if this mechanism was independent of TSC/mTOR, we next infected miR-126 knockdown and *tsc1a* knockdown embryos with *M*. *marinum* in the trunk region and measured the level of *tnfa* promoter activation around the site of infection using *TgBAC(tnfa:gfp)^pd1028^* ^44^. Expression of *tnfa* is a marker of classically activated bactericidal macrophages in zebrafish ^45^; we therefore utilised *tnfa* promoter activation as a surrogate marker of protective inflammatory, or classically activated, macrophages. At 1 dpi, miR-126 knockdown embryos had significantly reduced activation of the *tnfa* promoter, while knockdown of *tsc1a* alone increased *tnfa* promoter activation compared to control and miR-126 knockdown embryos (Figure 9C-D). Double knockdown rescued *tnfa* promoter activation to an intermediate level, indicating that *tsc1a*-independent mechanisms suppress *tnfa* promoter activation in miR-126 knockdown embryos.

Our data thus far indicates that miR-126 knockdown increases the recruitment of macrophages to *M*. *marinum* infection, and these macrophages are unable to clear bacteria, resulting in macrophage cell death. While the cell death phenotype can be attributed to increased *tsc1a* expression in miR-126 knockdown embryos, and *tsc1a* is partially responsible for the lack of *tnfa* promoter activation, there are clearly *tsc1a*-independent mechanisms responsible for the bulk recruitment of non-*tnfa* promoter active macrophages.

### Recruitment of permissive macrophages to *M*. *marinum* infection is dependent on Cxcl12a/Ccr2 signalling in miR-126 knockdown zebrafish embryos

We hypothesised that the increase in expression of Cxcl12/Ccl2/Ccr2 signalling axis components in miR-126 knockdown embryos is responsible for the recruitment of permissive macrophages to infection. To confirm the active role of the Cxcl12/Ccl2/Ccr2 signalling axis as a downstream effector mechanism of miR-126 expression and its involvement in the increased migration of permissive macrophages to sites of mycobacterial infection, the *cxcl12a* and *ccr2* genes were targeted for knockdown using Crispr-Cas9. Consistent with the work of Cambier *et al*., we did not observe any effect of *cxcl12a* or *ccr2* knockdown compared to control embryos across the three phenotypes of bacterial burden, total macrophage area, or *tnfa* promoter activity at sites of infection (Figure 10) ^46,47^.

**Figure 10.**
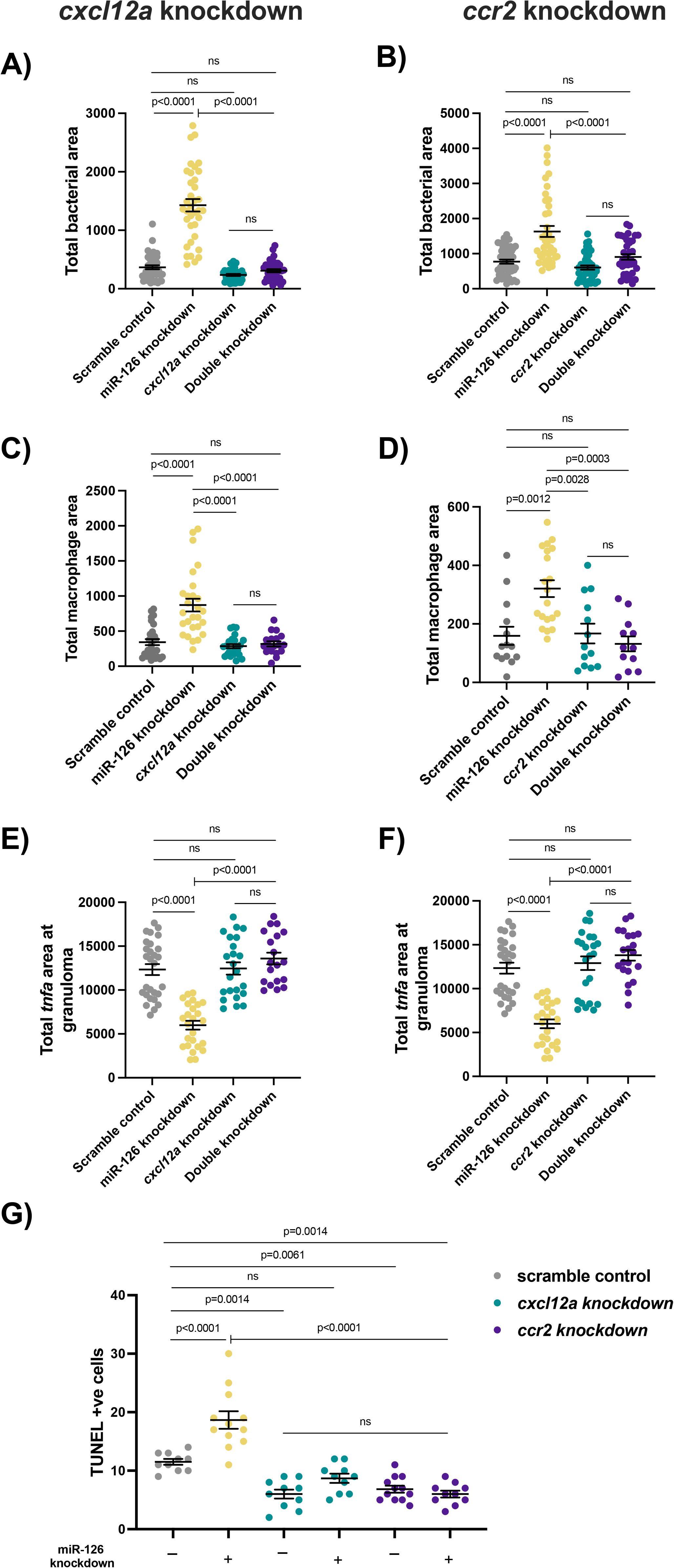
Infection-induced miR-126 regulates Cxcl12/Ccl2/Ccr2 signalling to prevent permissive macrophage recruitment to sites of infection. (A-B) Bacterial burden measured at 1 dpi in miR-126 and (A) *cxcl12a* or (B) *ccr2* knockdown embryos infected with *M*. *marinum*. (C-D) Whole-body macrophage levels measured at 1 dpi in miR-126 and (C) *cxcl12a* or (D) *ccr2* knockdown embryos infected with *M*. *marinum*. (E-F) *tnfa* fluorescent area at sites of infection measured at 1 dpi in miR-126 and (E) *cxcl12a* and (F) *ccr2* knockdown embryos infected *M*. *marinum*. (G) TUNEL +ve cells counted at 3 dpi in miR-126 and *cxcl12a* or *ccr2* knockdown embryos infected *M*. *marinum*. Each data point represents a single measurement, with the mean and SEM shown (n=10-45) embryos per group. Bacterial burden, macrophage analysis and granuloma *tnfa g*raphs are representative of 2 experimental replicates, while TUNEL staining is presented as a single replicate.

Knockdown of *cxcl12a* and double knockdown of both miR-126 and *cxcl12a* significantly reduced bacterial burden compared to miR-126 knockdown alone (Figure 10A). Similarly, knockdown of *ccr2* and double knockdown of both miR-126 and *ccr2* significantly reduced bacterial burden compared to miR-126 knockdown alone (Figure 10B). There was no difference in bacterial burden between double knockdown embryos and *cxcl12a* or *ccr2* knockdown alone embryos, suggesting the Cxcl12/Ccl2/Ccr2 signalling axis is driving the miR-126 knockdown-associated increase in burden.

Knockdown of *cxcl12a* and double knockdown of both miR-126 and *cxcl12a* significantly reduced total macrophage area compared to miR-126 knockdown alone (Figure 10C). Similarly, knockdown of *ccr2* and double knockdown of both miR-126 and *ccr2* significantly reduced total macrophage area compared to miR-126 knockdown alone (Figure 10D). Again, there was no difference in total macrophage area between double knockdown embryos and *cxcl12a* or *ccr2* knockdown alone embryos, suggesting the Cxcl12/Ccl2/Ccr2 signalling axis is driving the miR-126 knockdown-associated increase in macrophage numbers.

Following knockdown of *cxcl12a* and double knockdown of both miR-126 and *cxcl12a* restored *tnfa* promoter activation in granulomas to the levels in scramble control embryos (Figure 10E). Similarly, knockdown of *ccr2* and double knockdown of both miR-126 and *ccr2* restored *tnfa* promoter activation in granulomas to scramble control embryos (Figure 10F). As with the other phenotypes, there was no difference in *tnfa* promoter activation in granulomas between double knockdown embryos and *cxcl12a* or *ccr2* knockdown alone embryos.

Finally, we examined cell death in miR-126 and either *cxcl12a* or *ccr2* knockdown embryos to determine if the permissive nature of recruited macrophages further contributes to the increased cell death observed in miR-126 knockdown embryos (Figure 10G). As previously observed, miR-126 knockdown alone increased cell death compared to control infected embryos. Knockdown of down both *cxcl12a* and *ccr2* was protective, decreasing the number of TUNEL-stained cells at 3 dpi. Further, double knockdown of miR-126 and either *cxcl12a* or *ccr2* reduced cell death and from miR-126 knockdown alone, demonstrating that the Cxcl12/Ccl2/Ccr2 signalling axis is driving the miR-126 knockdown-associated increase in permissive macrophage recruitment to mycobacterial infection.

## Discussion

The ability of mycobacteria to evade host immunity is central to their pathogenicity and ability to establish a chronic infection. Control of miRNA is contested by host and invading mycobacteria as miRNA represent key nodes that are capable of exerting pleiotropic effects on the host immune response. Here, we show that increased abundance of miR-126 following infection with *M*. *marinum* is beneficial to the zebrafish host and prevents dissemination of bacteria during the early stages of infection. Through suppressing the target gene, *tsc1a*, and inhibiting the Cxcl12a/Ccl2/Ccr2 signalling pathway, miR-126 suppresses macrophage death and the recruitment of permissive macrophages. Our experiments demonstrate that decreasing miR-126 expression early during infection increases the recruitment of permissive macrophages which favours infection by failing to clear intracellular mycobacteria and facilitating mycobacterial release.

The expression of miR-126 was increased following infection with *M*. *marinum*, ΔESX1 *M*. *marinum*, and UPEC suggesting an early host-response to systemic infection. However knockdown of miR-126 increased the burden of WT *M*. *marinum*, not ΔESX1 *M*. *marinum* or UPEC. Thus, we considered the protective effect of miR-126 upregulation to be dependent on the mycobacterial ESX1 secretion system that is also present in *M*. *tuberculosis*, and this is a specific response to counteract the virulence of WT *M*. *marinum*.

The downstream effects of altered miR-126 expression were mediated in part, by the target gene *tsc1a*. As this gene negatively regulates mTORC1 activity, we were encouraged to find that the miR-126 knockdown phenotype was recapitulated by inhibition of mTOR and restored by co-knockdown of *tsc1a*. However, our data showing mTORC1 activity on distal to sites of infection, that inhibition of mTOR further exacerbated bacterial burden and cell death phenotypes in the miR-126 knockdown background, and that the small molecule activation of mTOR only partially rescued the miR-126 knockdown phenotypes suggest that while the miR-126/Tsc1a/mTOR axis is likely, it falls short of fully explaining the effects of miR-126 on the host response to *M*. *marinum* infection. While decreased phosphorylation of ribosomal protein S6 was observed in miR-126 knockdown embryos, indicating decreased mTOR activity, further experiments to assess the relative abundance of mTOR targets are required to provide conclusive evidence of a functional miR-126/Tsc1a/mTOR axis.

We observed that inhibition of mTOR with rapamycin is detrimental to the host, enhancing cell death and allowing for uncontrolled growth, whereas small molecule activation of mTOR is beneficial in the zebrafish-*M*. *marinum* infection model. Previous investigations into the role of mTOR in mycobacterial infection have uncovered a protective role for decreased mTOR signalling through improving mycobacterial killing ^48,49^, so that mTOR inhibitors, such as rapamycin, were considered potential host-directed therapies for the treatment of tuberculosis ^50^. While our results counter those observed in *in vitro* murine cell culture experiments, they are in agreement with observations from similar *M*. *marinum*-zebrafish models, where mTOR deficient zebrafish were more susceptible to mycobacterial infection, displaying severe disease and non-bactericidal macrophages^37^. This further reinforces the necessity of *in vivo* whole organism models of natural mycobacterial infection to capture the complex cellular interactions resulting from alterations of key signalling pathways.

We have previously identified a miRNA-mediated Cxcr4/Cxcl12 signalling axis which increased protective neutrophil responses in mycobacterial infection ^7^. Despite an increase in expression of these same genes following knockdown of miR-126, we found no change in neutrophil recruitment to infection in the current study. This led us to investigate alternative receptors for the Cxcl12 ligand in our miR-126 knockdown embryos that may contribute to the enhanced macrophage influx. Mammalian CXCL12 can form heterodimers with CCL2 to bind to the CCR2 receptor ^51^.

Pathogenic mycobacteria utilise membrane lipids to recruit non-bactericidal Ccr2+ permissive macrophages and ensure bacterial persistence ^46,47,52^. However, this observation of Ccr2-dependant macrophage recruitment has only been previously observed in zebrafish infected with *M*. *marinum* in the hindbrain protected by the blood-brain barrier, and not in systemic infection despite the upregulation of *ccl2* after infection of either site ^47^. In this study we observed increased bacterial burden associated with the lack of *tnfa:gfp* positive inflammatory macrophages in the contexts of systemic infection and at localised sites of infection in the trunk musculature only when miR-126 was depleted. The miR-126 knockdown-induced increase in *ccr2* transcription was independent of infection suggesting miR-126 is required for the physiological suppression of Ccr2+ macrophage differentiation, thus the pool of available macrophages in our miR-126 knockdown embryos is skewed towards permissive macrophages even before infection.

While we have associated miR-126 with *tsc1* and the *cxcl12a/ccl2/ccr2* axis by combinatorial gene depletion studies, miR-126 has been documented to be involved in additional pathways that may impact the outcome of infection. miR-126 has well characterised roles in the formation of blood vessels and lymphatics ^53–57^. Angiogenesis is a major host pathway appropriated by pathogenic mycobacteria to promote pro-angiogenic programmes and increase vascular permeability, enabling bacterial dissemination ^58,59^. It is therefore possible that although increased miR-126 is host protective through preventing permissive macrophage accumulation, pathogenic mycobacteria may co-opt this pathway to increase infection-induced angiogenesis around mature granulomas after the time period studied in our present work.

Another identified target gene of miR-126 which may be active in mycobacterial infection is *spred1*, and this could further contribute to angiogenic signalling through a Spred1/Vegf axis ^60,61^. Beyond the recognised angiogenic role, Spred1 has been associated with host immunity through regulation of haematopoietic homeostasis, mast cell activation, and eosinophil infiltration ^62–64^. The involvement of both mast cell- and eosinophil-related bactericidal mechanisms during mycobacterial infection has been established, and the mediation of these cellular responses by upstream miR-126 may further contribute to the host-protective effect of infection-induced miR-126 ^65–67^. It is evident that miR-126 has numerous target genes involved in a variety of cellular pathways relevant to mycobacterial infection, and that its effect on angiogenesis during disease certainly warrants further investigation in a model with established mycobacterial granulomas.

miR-126 has previously been identified as having decreased expression in both plasma from pulmonary tuberculosis patients, and PBMCs of tuberculous meningitis patients ^13,16^. We have observed that a reduction in miR-126 transcript alters normal macrophage phenotype and function through its involvement in the Cxcl12/Ccl2/Ccr2 axis, and coupled with concurrent mTOR inhibition, increases death of permissive macrophages. The lack of bacterial containment and failure to control infection seen in our miR-126 knockdown embryos may provide an insight into the effect of reduced miR-126 transcript abundance in *M*. *tuberculosis* infection. As macrophages are a primary intracellular niche for pathogenic mycobacteria, the subversion and disruption of their normally protective function towards a permissive state by miRNA adds another level to their complex regulatory networks. Identification of these networks and the molecules targeted by pathogenic mycobacteria may provide new avenues for host-directed therapies to prevent progression to chronic disease.

In this study we have identified several mechanistic functions of miR-126 in mycobacterial infection. The analysis of potential target genes revealed a link between altered miR-126 expression and the dysregulation of host macrophage responses. We demonstrate that infection-induced miR-126 suppresses *tsc1* and *cxcl12a* expression thus improving macrophage function during the early stages of infection, partially through activation of mTOR signalling and strongly through preventing the recruitment of Ccr2+ permissive macrophages. The increased macrophage responses are likely to be mediated by a combination of Tsc1/mTOR suppression of cell death, enhanced activation of bactericidal macrophages, and suppression of permissive macrophage recruitment. Through utilising a zebrafish-*M*. *marinum* model, we were able to identify complex multicellular interactions from converging biological pathways that alter the course of mycobacterial infection. These responses appear to be conserved across mycobacterial infection in vertebrate hosts and provide further insight into the intricate regulation of immunity by miRNA.

## Methods

### Zebrafish husbandry

Adult zebrafish were housed at the Centenary Institute and breeding was approved by Sydney Local Health District AWC Approval 17-036. Embryos were obtained by natural spawning and were raised in E3 media and maintained at 28-32°C.

### Zebrafish lines

Zebrafish were AB strain. Transgenic lines used were: *Tg(lyzC:GFP)^nz117^* and *Tg(lyzC:DsRed2)^nz50^* for neutrophil imaging experiments ^68^, and *Tg(mfap4:turquoise)^xt27^* and *Tg(mfap4:tdtomato)^xt12^* for macrophage imaging experiments ^69^.

### Embryo microinjection with antagomiR

Embryos were obtained by natural spawning and were injected with either miR-126 antagomiR (-GCAUUAUUACUCACGGUACGA-) or a scramble control (-CAGUACUUUUGUGUAGUACAA-) (GenePharma, China) at 200 pg/embryo at the single cell stage and maintained at 32°C.

### miRNA target prediction

Prediction of target mRNA was performed using TargetScan. dre-miR-126a-3p was entered into TargetScanFish 6.2 (http://www.targetscan.org/fish_62/), hsa-miR-126-3p entered into TargetScan 7.2 (http://www.targetscan.org/vert_72/), and mmu-miR-126a-3p entered into TargetScanMouse 7.2 (http://www.targetscan.org/mmu_72/). Experimentally validated targets were compiled using miRTarBase (release 8.0) (http://mirtarbase.cuhk.edu.cn). Prediction of binding of miR-126 and *tsc1a* in zebrafish was performed using RNAhybrid (https://bibiserv.cebitec.uni-bielefeld.de/rnahybrid) ^70^.

### *M*. *marinum* culture

*M*. *marinum* was cultured as previously described ^71^. Briefly, *M*. *marinum* M strains expressing Wasabi, Katushka, or tdTomato fluorescent protein were grown at 28°C in 7H9 supplemented with OADC and 50 μg/mL hygromycin to an OD600 of approximately 0.6 before being washed and sheared by aspiration through a 32 G needle into single cell preparations. These were then aliquoted and frozen in 7H9 at −80°C until needed. The concentration of bacteria was quantified from thawed aliquots by CFU recovery onto 7H10 supplemented with OADC and 50 μg/mL hygromycin and grown at 28°C.

### UPEC culture

Uropathogenic *Escherichia coli* (UPEC) carrying the mCherry PGI6 plasmid ^72,73^ was cultured as previously described ^7,74^. Briefly, bacteria were cultured in LB supplemented with 50 μg/mL of spectinomycin overnight at 37°C with shaking at 200 rpm. Bacteria were then further diluted 1:10 with LB + spectinomycin (50 μg/ml) and incubated for 3 hours at 37°C with 200 RPM shaking. Culture medium/broth (1 ml) was centrifuged (16,000 × *g* for 1 minute), and the pellet washed in PBS. The bacterial pellet was resuspended in 300 μl of PBS + 10% glycerol and aliquoted for storage. Enumeration of bacteria was performed by serial dilution on LB + spectinomycin agar plates and culturing at 37°C overnight. Bacterial concentration was determined by CFU counts.

### Bacterial infections

Embryos were dechorionated and anesthetised in tricaine (160 μg/ml) and staged at approximately 1.5 dpf. Working solutions of *M*. *marinum* or UPEC (diluted with 0.5% w/v phenol red dye) were injected into either the caudal vein or trunk to deliver approximately 200 CFU *M*. *marinum* or 250 CFU UPEC. Embryos were recovered in E3 media + PTU (0.036 g/L) and maintained at 28°C.

### Crispr-Cas9 mediated knockdown

Embryos were injected at the 1-2 cell stage with 1 nl of Crispr mixture containing 1 μg/μl gRNA, 500 μg/mL Cas9. For double knockdowns with Crispr-Cas9 and antagomiR, mixtures contained 1 μg/μl gRNA, 100 pg/nl antagomiR, 500 μg/mL Cas9. gRNA was synthesised as previously described (47). Embryos were transferred to E3 containing methylene blue and maintained at 32°C.

### Gene expression analysis

Groups of 10 embryos were lysed and homogenised using a 27-gauge needle in 500 μl Trizol (Invitrogen) for RNA extraction. cDNA was synthesised from 500 ng RNA using the miScript II RT kit with HiFlex buffer. qPCR was carried out on an Mx3000p Real-time PCR system using Quantitect SYBR Green PCR Mastermix and primer concentration of 300 nM (Table 2.). For miRNA qPCRs, the miScript Universal Primer was used alongside miR specific miScript primer assays (miR-126 GeneGlobe ID MS00005999 and U6 cat. no. MS00033740).

**Table 1.**
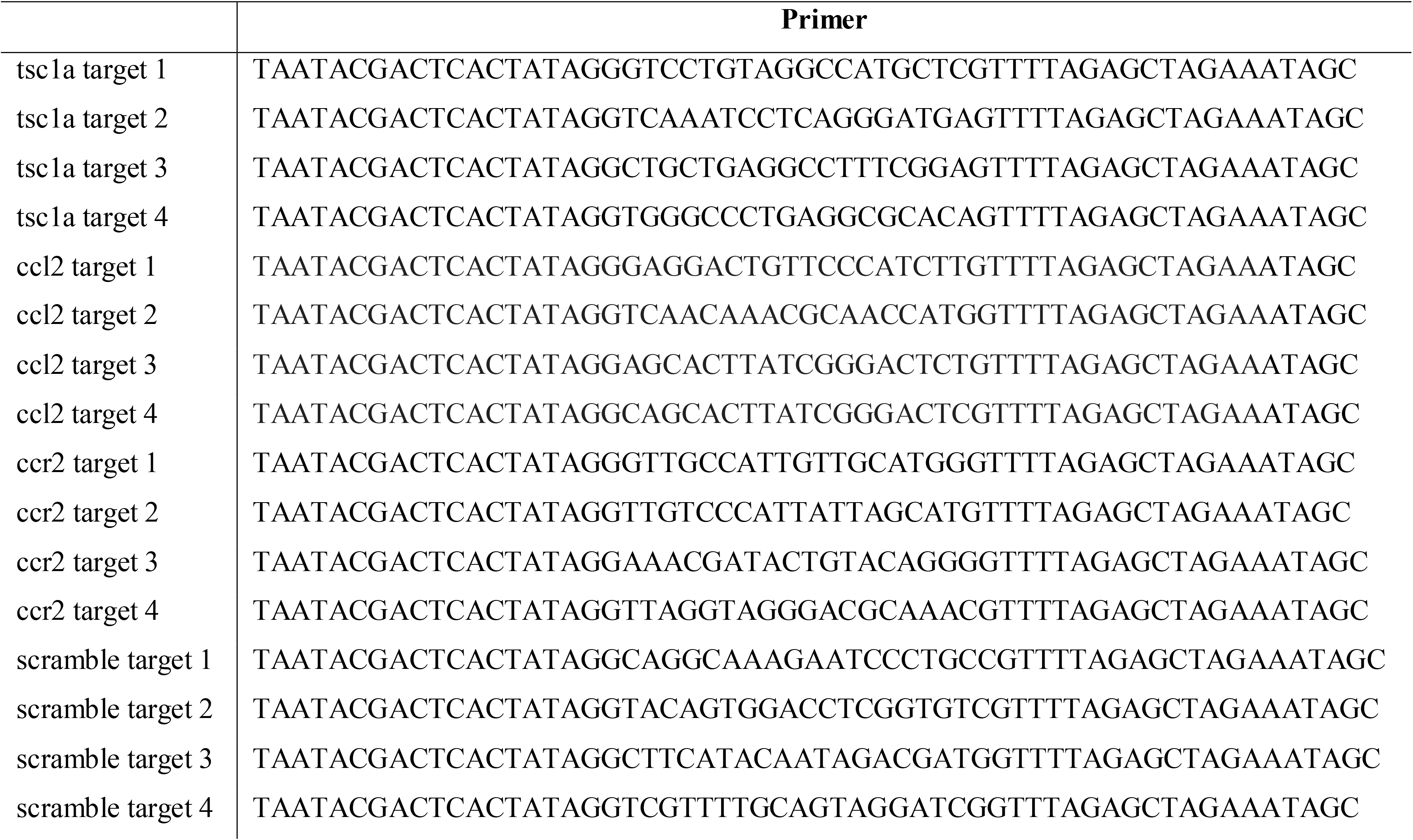
Guide RNA sequences used for Crispr-Cas9 mediated knockdown experiments.

**Table 2.**
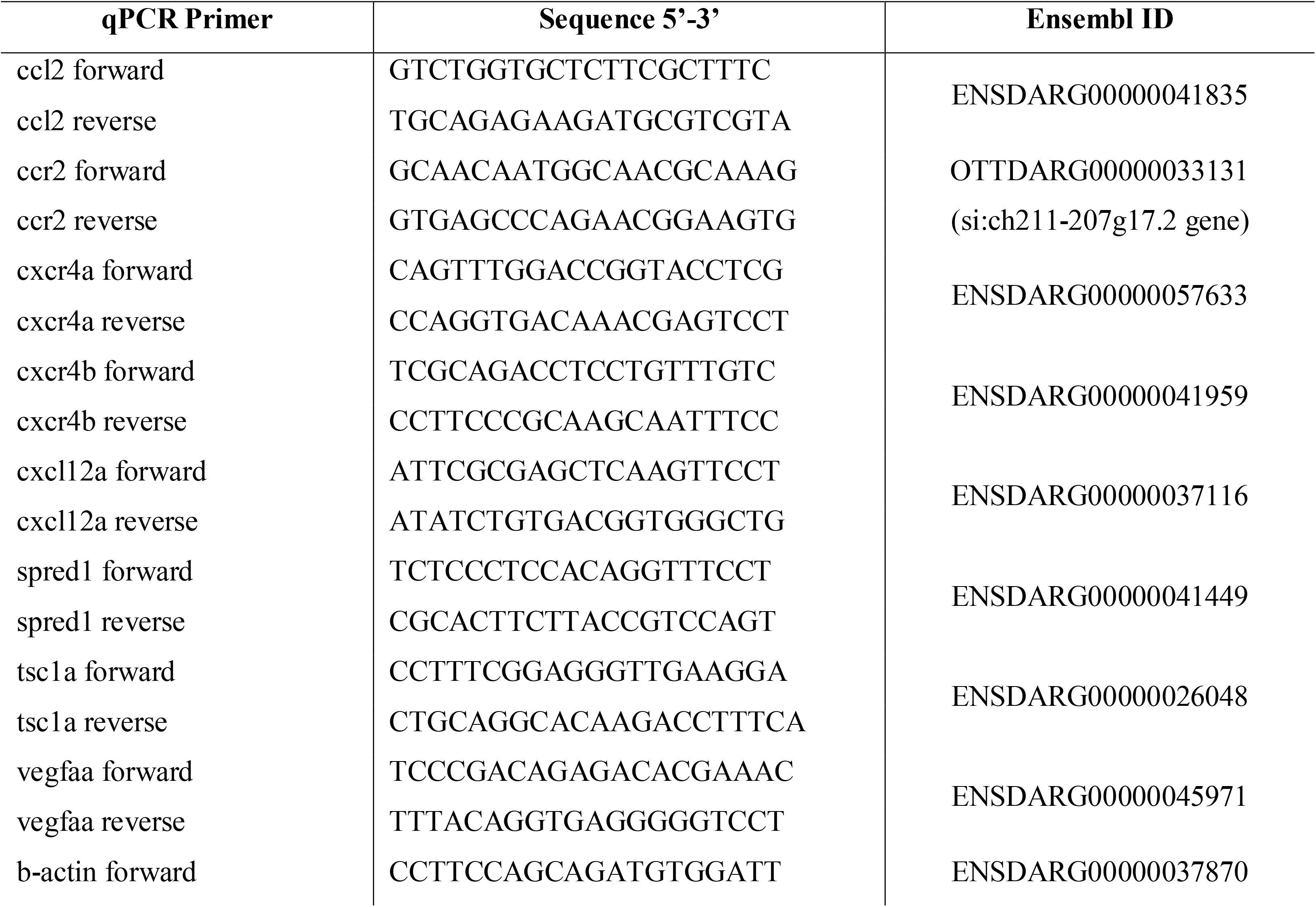
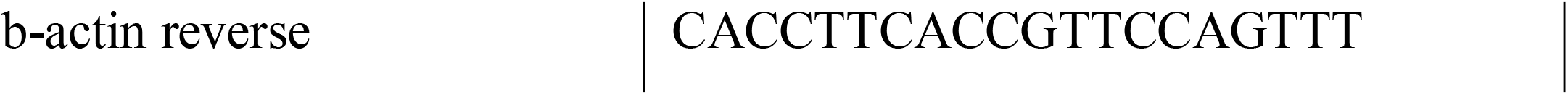
qPCR primer sequences.

Cycling conditions for miRNA were: 95°C for 15 minutes; 40 cycles of 95°C for 20 seconds, 56°C for 30 seconds, 72°C for 30 seconds with fluorescence data acquisition occurring at the end of each cycle, followed by 1 cycle of 95°C for 1 minute, 65°C for 30 seconds, and 97°C for 30 seconds. For mRNA, conditions were: 95°C for 15 minutes; 40 cycles of 94°C for 15 seconds, 55°C for 30 seconds, 70°C for 30 seconds with fluorescence data acquisition occurring at the end of each cycle, followed by 1 cycle of 95°C for 1 minute, 65°C for 30 seconds, and 97°C for 30 seconds.

U6 or β-actin was used as an endogenous control for normalisation and data analysed using the 2^−ΔΔ^ Ct method.

### Drug treatment

Embryos were treated with either 50 nM rapamycin (Sigma-Aldrich, USA) or 5 μm MHY1485 (Sigma-Aldrich, USA) dissolved in DMSO and refreshed daily.

### Static imaging and burden analyses

Live imaging was performed on anaesthetised embryos on a depression microscope slide. Images were acquired using a Leica M205FA Fluorescent Stereo Microscope equipped with a Leica DFC365FX monochrome digital camera (Leica Microsystems, Germany). Images were analysed using ImageJ software to quantify the fluorescent pixel count, defined as fluorescent signal above a consistent set background determined empirically for each experimental dataset ^71^. Data are presented as total fluorescent area (pixels) above background level.

For static imaging of granuloma associated macrophages, cells within a 200 μm box surrounding bacterial granulomas were measured and classified as “granuloma-associated macrophages”. Expression of GFP in the *tnfa:gfp* line was measured within a 500 μm box around infection foci.

### Macrophage and neutrophil tracking analyses

Time-lapse imaging was performed on a DeltaVision Elite at 28°C (GE, USA). Following infection with *M*. *marinum* into the trunk, embryos were mounted in a 96-well black-walled microplate in 1% low-melting point agarose topped up with E3. Images were captured every 60-180 seconds for 16-24 hours. Analysis was performed using ImageJ software. Briefly, every 10-30 images were analysed for the quantity of neutrophils or macrophages in a 1000 x 500 μm box around infection foci by quantifying the fluorescent pixel count (total area) at each time point.

### TUNEL cell death staining

Embryos were infected with *M*. *marinum* via caudal vein injection and analysed for apoptotic cells at 1- and 3- dpi using the Click-iT™ Plus TUNEL Assay (Thermo Fisher, USA) according to the manufacturers protocol. Briefly, embryos were fixed in 10% neutral buffered formalin (NBF) overnight at 4°C. Embryos were permeabilised with proteinase K (10 μg/mL) for 30 minutes at room temperature and re-fixed in 10% NBF. edUTP and Alexa Fluor™ 488 incorporation reactions were performed at 37°C protected from light. Embryos were imaged on a DeltaVision Elite (GE, USA) and TUNEL stained cells counted using the Multi-point tool in ImageJ.

### Embryo whole-mount immunofluorescence

Embryos were infected with *M*. *marinum* via caudal vein injection and stained for phosphorylated ribosomal protein S6. At 1 dpi, embryos were fixed in 4% paraformaldehyde (PFA) overnight at 4°C followed by several washes with PBS + Tween 20 (PBST), and permeabilisation with proteinase K (10 μg/mL) for 15 minutes. Embryos were then washed with PBST and refixed in 4% PFA for 20 minutes. Samples were blocked in 5% goat serum prior to incubation with the primary antibody (P-S6 Ser235/236 1:100, 4856S Cell Signalling Technologies, USA) overnight at 4°C with gentle rocking. Embryos were then washed with PBST and blocked with 5% goat serum before addition of the secondary antibody (Goat anti-rabbit IgG H&L DyLight 650, 1:200, 84546 Invitrogen, USA) and incubated overnight at 4°C with gentle rocking. Embryos were thoroughly washed with PBST and transferred to a 1:1 PBS/glycerol solution prior to imaging. Embryos were imaged on a DM600B (Leica Microsystems, Germany), and phospho-S6 quantified using the fluorescent pixel count (total area).

### Statistics

Statistical analysis was performed in GraphPad Prism (v. 9.0.0). All data were analysed by t-test or ANOVA depending on the experimental design, and comparisons between groups performed using Tukey’s multiple comparisons test. For time-lapse data, group comparisons were computed using the Sidak test. Outliers were removed prior to statistical analysis using ROUT, with Q=1%.

## Acknowledgements

The authors would like to acknowledge and thank Dr Angela Kurz of the BioImaging Facility and Sydney Cytometry at Centenary Institute for technical assistance with imaging, Drs Pradeep Manuneedhi Cholan and Kaiming Luo for assistance with imaging, and Dr Elinor Hortle for assistance with image analysis. We would like to acknowledge Ms Lina Daniel and A/Prof Carl Feng for providing reagents. We would also like to thank Professor Lalita Ramakrishnan and Dr Antonio Pagan for valuable insights.

## Author contributions

KW designed and performed experiments, wrote draft manuscript, secured funding. KdS, KMP, and ACP secured funding, edited manuscript. WJB secured funding, edited manuscript. SHO designed experiments, secured funding, wrote draft manuscript.

## Supplementary File legends

**Supplementary File 1**

Possible gene targets of miR-126 from database sources as annotated.

**Supplementary Figure 1**

Expression of *ccl2* as measured by RT-qPCR in (A) uninfected and (B) *M*. *marinum*-infected *cxcl12a* and miR-126 knockdown embryos.

**Supplementary Video 1**

Time lapse imaging of macrophage (magenta) recruitment to *M*. *marinum* (green) in *Tg(mfap4:turquoise)* control embryo.

**Supplementary Video 2**

Time lapse imaging of macrophage (magenta) recruitment to *M*. *marinum* (green) in *Tg(mfap4:turquoise)* miR-126 knockdown embryo.

**Supplementary Video 3**

Time lapse imaging of macrophage (magenta) recruitment to a tail wound on the right edge of the field of view in *Tg(mfap4:turquoise)* control embryo.

**Supplementary Video 4**

Time lapse imaging of macrophage (magenta) recruitment to a tail wound on the right edge of the field of view in *Tg(mfap4:turquoise)* miR-126 knockdown embryo.

