## Supplementary figures and images for "Infection-induced miR-126 suppresses *tsc1*- and *cxcl12a*-dependent permissive macrophages during mycobacterial infection"

### Supplementary Figure 1

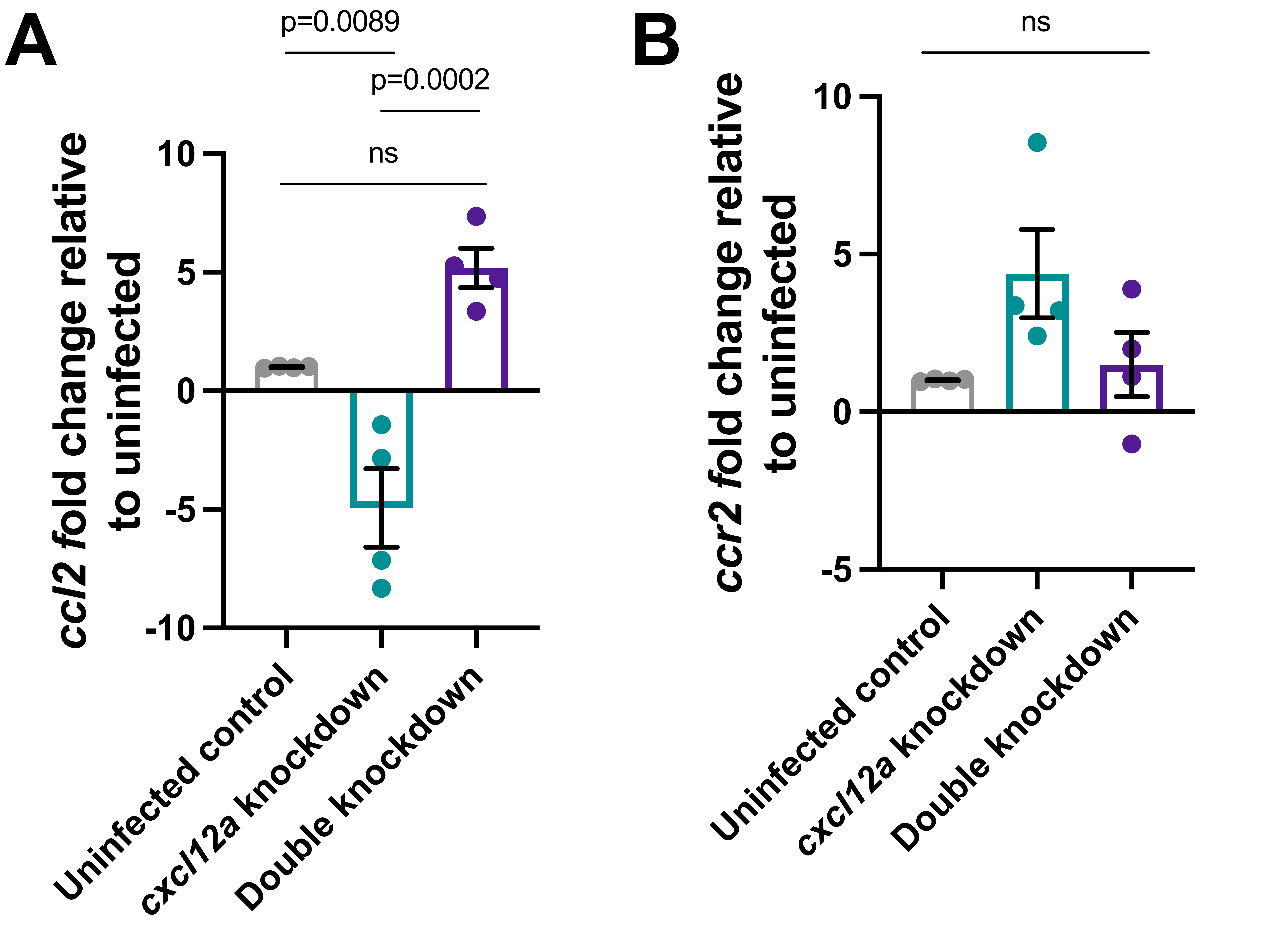
